# Genetic backgrounds and sex modulate indirect genetic effects of *Epha4* in laboratory mice

**DOI:** 10.64898/2026.09.30.755759

**Authors:** Pavan Matani, Micaela Stedile, A. Silvina Nacht, Amelie Baud

## Abstract

The phenotype of a focal individual can be affected not only by the individual’s own genes (direct genetic effects, DGE) but also by genes of other individuals it interacts with (indirect genetic effects, IGE). Better understanding IGE can help better understand the impact and mechanisms of social effects. Here, we investigated, in laboratory mice, the modulation of IGE by genetic background of the focal individual (“social epistasis”) and by sex. We focused on IGE arising from *Epha4*, a gene that has been associated with the individual’s own stress-related and social behaviours (i.e. DGE) and the stress-coping behaviour and wound healing of the other mice in the cage (i.e. IGE). Among four different genetic backgrounds for focal mice, significant IGE from *Epha4* were detected in CBA females only. A formal model selection analysis was consistent with the modulation of IGE by the similarity of the genetic backgrounds of the focal mouse and its Epha4 cage mate, as well as by sex. Observations of fighting in cages of males with dissimilar genetic backgrounds suggested aggression might underpin these modulatory effects, which prompted to us to expand the conceptual framework describing the mechanisms of social epistasis. Thus, our results strengthen the emerging view that IGE are not fixed properties of social genotypes but can depend on the genetic and sex context of interacting individuals, revealing an underexplored layer of their genetic architecture.

**Author summary:** The phenotype of a focal individual can be affected by genes of other individuals it interacts with (“indirect genetic effects”, IGE). The extent to which IGE depend on the context, in particular the genetic background and sex of the interacting individuals, is poorly understood. This study focused on IGE from *Epha4* in male and female laboratory mice and measured stress-coping behaviour and ear wound healing in focal mice from four genetic backgrounds co-housed with a cage mate with or without a mutation in *Epha4*. The effects of partner *Epha4* genotype varied across genetic backgrounds and sexes. We expanded the conceptual framework describing how the context can modulate IGE. Our results have important implications for the design of IGE experiments.

## Introduction

Conspecifics are an important component of the environment in social species as traits they express can influence the phenotype of a focal animal. When the influential traits of social partners have a genetic basis, “indirect genetic effects” (IGE, also called social genetic effects) arise from the genotypes of social partners (1–4). These IGE are in addition to “direct genetic effects” (DGE), which arise from the focal individual’s own genotypes.

IGE have been detected in a large number of species, contexts (e.g. parents/offspring, peers) and phenotypes (5,6). Characterising the mechanisms of IGE, and in particular identifying the heritable traits of social partners giving rise to IGE, is now recognised as the most pressing challenge across all disciplines concerned with IGE (biomedical genetics, animal and plant breeding, conservation science, etc.) (4). The challenge is greatest when the goal is to discover traits of social partners mediating IGE in the absence of any prior hypothesis, an approach that is necessary to gain an unbiased view of the mechanisms of IGE. IgeGWAS is a promising approach to do so (7–14). A similar strategy has been successfully used with classical (DGE) GWAS to identify biological pathways within the individual that play a role in the phenotype (15).

The identification of an IGE locus affecting stress-coping behaviour (measured as immobility in the Forced Swim Test (FST)) and ear wound healing mapping to a topological domain encompassing a single gene, *Epha4*, in outbred Swiss Webster (CFW) laboratory mice, and the subsequent detection of IGE in inbred FVB mice paired with a wild-type (*Epha4^+/+^*) or heterozygous knock-out (*Epha4^+/^*) cage mate (14), provide an unprecedented opportunity to leverage an igeGWAS result and to evaluate this approach. However, Baud et al. noted clear differences between the magnitude of IGE detected in CFW and FVB mice, and subtler differences between same-sex groups of males and females (Table 1), suggesting that the genetic background of the focal (phenotyped) mice and, perhaps also, the sex of the co-housed mice modulated IGE.

**Table 1.** Evidence of strain- and sex-dependent IGE from *Epha4* affecting immobility in the FST (a measure of stress-coping behaviour) and ear hole area (a measure of wound healing) in laboratory mice. Summary of results from Baud et al. (**14**).

| Focal mice | Analysis | Phenotype | Result |
| --- | --- | --- | --- |
| CFW | igeGWAS for discovery (males and females analysed jointly) | Immobility first 2 min. FST | genome-wide significant (-logP = 5.2) |
|  |  | Immobility last 4 min. FST | genome-wide significant (-logP = 2.8) |
|  |  | Ear hole area | Suggestive but not genome-wide significant |
| FVB | ANOVA comparing pairing with an *Epha4*<sup>+/+</sup> or *Epha4*<sup>+/-</sup> cage mate | Immobility first 2 min. FST | P > 0.05 in males and females (immobility too low - floor effect) |
|  |  | Immobility last 4 min. FST | p = 0.054 in males; p > 0.05 in females |
|  |  | Ear hole area | p = 0.038 in males; p = 0.05 in females |
| CFW | retrospective comparison of effect sizes in males and in females | Immobility first 2 min. FST | Not analysed due to FVB results |
|  |  | Immobility last 4 min. FST | effect size in males similar to in females |
|  |  | Ear hole area | effect size in males greater than effect in females |

The genetic background is known to often modify the phenotypic consequences of genetic mutations. This was most clearly demonstrated by knocking out, in laboratory mice, the genes *Cacnac1* or *Tcf7l2* in 30 different genetic backgrounds. The same mutation increased, decreased or did not change the measured behaviour, depending on the background (16). Similarly, the genotypes of focal individuals can modulate the magnitude of IGE, as first described by Wolf (17) (genotype-by-genotype or among-genotype epistasis, also referred to as GxG epistasis, GxG IGE, DGExIGE interactions (4), and intergenomic social epistasis (18), which we choose to simplify to “social epistasis”). The theoretical and empirical literature relevant to social epistasis is fragmented across the various modelling approaches traditionally used to study IGE (phenotype-first frameworks (19,20), trait-based models (6), and variance components models (21)), the type of genetic variation (single genetic variants/loci (17,22), genetically-determined color morphs(23), inbred strains (18,24), and outbred populations (21)) and how social epistasis arises (dependent on the genotypic values of the interacting individuals (25) or on the similarity between their genotypes (23,24). A number of studies on same-genotype vs. mixed-genotype housing of laboratory strains is also relevant here, despite not mentioning IGE and not framing the results as social epistasis (22,26). Similarity-dependent social epistasis is expected from kin-selection theory, which predicts that individuals behave differently towards kin, who are genetically similar, as compared to strangers (21) and in settings where the social environment sets norms for behavior (27), and it has previously been observed in experimental systems.

A modulating effect of sex has also been reported for both DGE and, to a lesser extent, IGE (7,28,29).

Given prior evidence of context-dependent IGE from *Epha4*, the goals of this study were (i) to identify a suitable experimental system (strain and sex) where IGE from *Epha4* can be robustly detected, so that, in the future, it can be used for identifying the trait(s) of cage mates mediating IGE from *Epha4*; (ii) to further characterise the context-dependency of IGE. Focal (phenotyped) mice were from four different inbred genetic backgrounds (C57BL/6J “B6”, CBA/J “CBA”, DBA/2J “DBA” and MRL/MpJ “MRL”, Fig. 1A), chosen because they differed greatly in the phenotypes studied, stress-coping behaviour and ear wound healing (30–33). As we wanted to identify a focal strain in which IGE can be robustly detected, we determined *a priori* the sample size necessary for detection of IGE within each strain, based on the effect size estimated in FVB mice (N = 98), resulting in a large-scale experiment with about 400 focal mice and 400 cage mates. Secondly, we formally evaluated whether focal genetic background and sex modulated IGE from *Epha4*, using the whole dataset (Fig. 1B). To avoid potential batch effects confounding the latter analysis, all four focal strains were tested concurrently using the same protocols.

**Fig 1.**
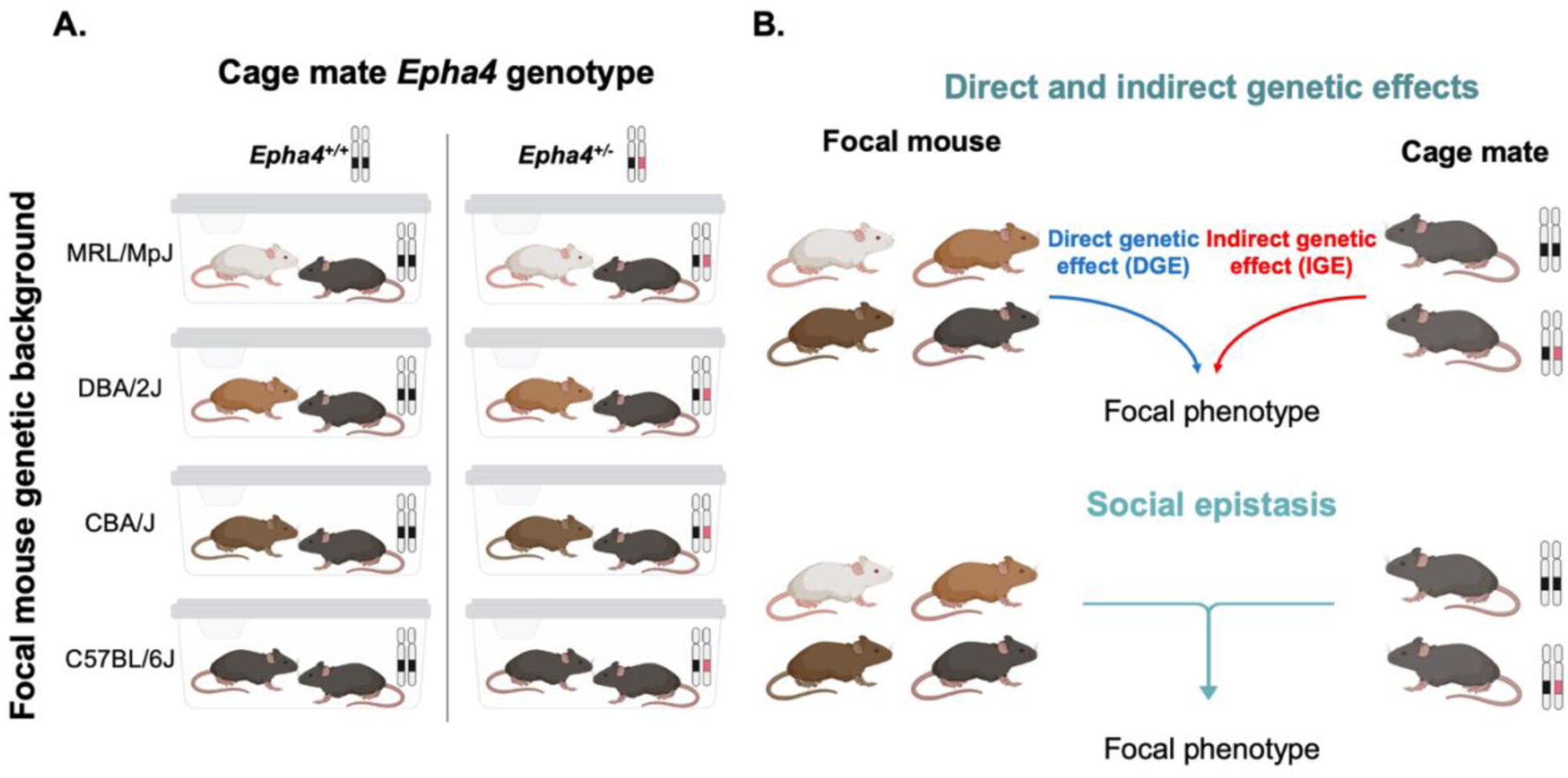
Co-housing of focal mice and cage mates and definition of social epistasis. **A.** Co-housing of focal mice from four strains (MRL/MpJ “MRL”, DBA/2J “DBA”, CBA/J “CBA” and C57BL/6J “B6” and, on the left in each cage) with cage mates that varied in their *Epha4* genotype (on the right in each cage). Sample size for every such combination was approximately 50. The genetic background of the cage mates was the same as the genetic background of one of the focal strains (B6), hence the same dark grey colour. **B.** Definition of social epistasis. In a simple scenario, the genetic background of the focal animal and the *Epha4* genotype of the cage mate independently affect the focal phenotype (DGE and IGE acting additively). In contrast, social epistasis occurs when the genetic background of the focal animal modulates the effect of the *Epha4* genotype of the cage mate (i.e. a statistical interaction between DGE and IGE). That modulation could depend either on the strain of the focal mouse (“values-dependent epistasis”, see main text), or on the similarity between the genetic backgrounds of the two mice in the cage (“similarity-dependent epistasis”). The genetic background of the Epha4 cage mates is B6 (see Methods), hence cages with a B6 focal mouse represent cages with same genetic backgrounds, whereas cages with CBA, DBA or MRL focal mice represent mixed genetic backgrounds. Modulation of IGE by same VS mixed genetic backgrounds and B6-specific modulation of IGE are not distinguishable in our design, given that all social partners (Epha4 line) have a B6 background.

Analysis of large-scale behaviour experiments such as this one, in which just under 400 animals are used, is labor-intensive and essentially impossible without an automated or semi-automated method for characterising mouse behaviour. A few such methods have been developed to quantify immobility in the FST (34–40). The methods based on computer-vision and deep learning perform well (34,37) but they cannot be used in new experimental setups without retraining the underlying computational models. In contrast, the methods based on key-point detection (e.g. nose, paws, base and tip of the tail), for example by the widely used software EthoVision (39,40), are not ideal because on a projection of the horizontal surface where the mouse swims some key-points may not move when the mouse is swimming, and conversely the movement of key-points is not a reliable indicator of swimming (e.g. the mouse can be drifting, or just moving a paw to stay afloat). To overcome these issues, we developed a new computational pipeline for semi-automated analysis of FST immobility that requires minimal adaptation across experimental conditions and that leverages full body movement information extracted from EthoVision.

## Results

### A pipeline for semi-automated analysis of forced swim test videos for characterising stress-coping behaviour

A computational pipeline was developed to quantify immobility during the first two and last four minutes of the FST, two measures widely used to study stress-coping behaviour in laboratory rodents (see Methods and Figure 2). Briefly, the animal tracking system EthoVision XT (41) was used to detect movement through changes in pixel intensity across frames. Then, strain-specific thresholds for converting pixel-intensity data to a binary call (mobile/immobile) for each frame were determined. This was done by manually annotating all six minutes of a small subset of videos (training set) and maximizing the F-1 score, which is a compound measure of accuracy and recall calculated, for each video, from the frame-level comparison of manual and automatic immobility calls (Supplementary material, Figure S1). Finally, the whole pipeline was evaluated using an independent set of videos (test set) that were also manually annotated and compared to automatic predictions using both the F-1 score, which is widely used by the machine learning community, and the Pearson correlation between manually derived and automatically derived immobility time, which is more commonly used by the animal behaviour community. The average F-1 score was 0.89 (a value of 1 means perfect agreement with the reference, here the manually annotated immobility calls) and the Pearson correlation 0.78 (Figure2D).

**Figure 2.**
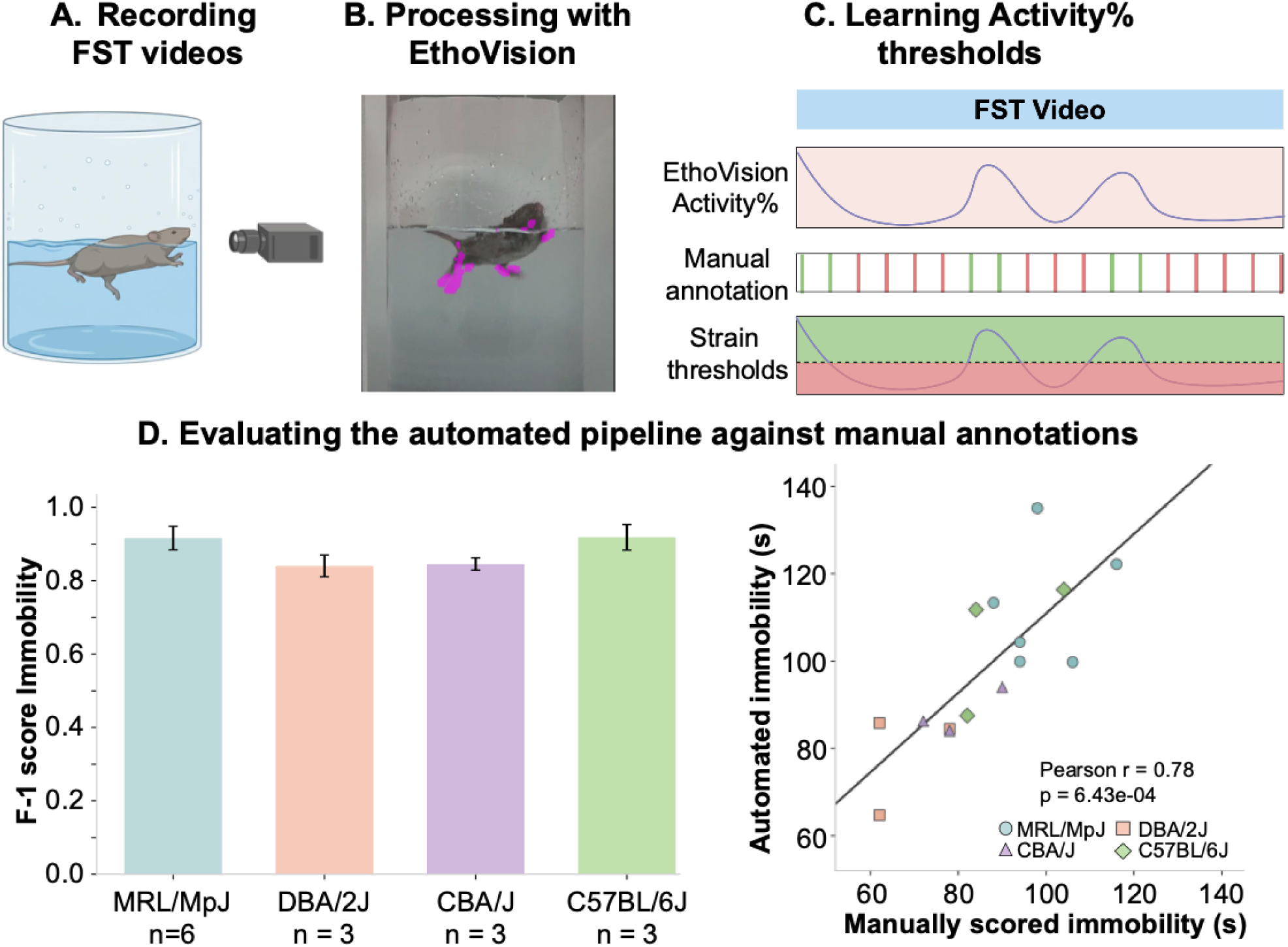
Forced Swim Test video recording, analysis method and validation. **A**. Video-recording set-up for the forced swim test (FST). **B.** Processing of the videos with EthoVision XT to obtain a per-frame measure activity (“Activity%”). **C.** Transformation of Activity% into per-frame mobile/immobile calls, based on strain-specific thresholds determined by comparing automated and human-made mobile/immobile calls in a subset of videos (training set). **D.** Evaluation of the pipeline using a separate subset of videos (test set), using F-1 scores (error bars indicate standard error of the mean) and the Pearson correlation between pipeline- and human-derived immobility times. MRL validation was separate for two batches of videos with different lighting, resulting in twice as many videos being used compared to the other strains. See Methods for full details. Figure created with BioRender.

To illustrate the ability of our pipeline to detect systematic biological differences, we compared the immobility time of the four focal strains, as these strains are known to exhibit different stress-coping behaviours (42–45). For this specific analysis, we used only the animals that were housed with wild-type *Epha4^+/+^* cage mates. Immobility time differed significantly across strains in both the first two minutes of the FST (One-way ANOVA: F (3, 204) = 44, p = 4.4e-22) and the last four minutes (One-way ANOVA; F = 44, p = 3.7e-22) (Supplementary material, Figure S2). The strain pattern that we observed, specifically higher immobility of B6 mice compared to CBA and DBA mice, has been reported in the literature (42–45). The MRL strain had not been compared to any of the other three strains before.

### Per-strain analysis of IGE from *Epha4*

Our experiment was designed with a per-strain analysis in mind, given that prior results clearly suggested the genetic background of the focal mice might modulate IGE from *Epha4* (Table 1) (14). In the power analysis conducted prior to starting this experiment to determine per-strain sample size, we used the effect size estimated for immobility during the last four minutes of the FST in FVB males and assumed similar effect sizes in males and females, based on inconsistent sex differences between CFW and FVB mice (Table 1 and see Methods). Thus, we first tested for IGE from *Epha4* in each focal strain separately considering males and females together (14). Doing so, no IGE from *Epha4* were significant (analysis of variance - ANOVA p-value > 0.05 for all four focal strains). (Supplementary material, Table S2).

To formally evaluate the possibility of sex-specific IGE from *Epha4*, we carried out a model comparison analysis using the Akaike Information Criterion (AIC). The results showed that the model including an interaction between sex and *Epha4* genotype of the cage mate was the best model for CBA mice for both FST phenotypes, all other models being noticeably worse (ΔAIC > 2 (46), Supplementary material, Tables S3-5). Thus, evidence of sex-dependent IGE on FST phenotypes in CBA was strong. In contrast, the interaction model was the best model for immobility during the first two minutes of the FST in B6 mice but the null model without sex or genotype was competitive (ΔAIC < 2, Supplementary material, Tables S3-5). Similarly, the intersection model was among the competitive models (but not the best) for immobility during the first two minutes of the FST in DBA mice, and for wound healing in MRL and DBA mice. In those cases, too small sample sizes might have limited the ability to discriminate among the competitive models, including the model with an interaction. Finally, for all other phenotype - strain combinations, the interaction model was not among the competitive models (ΔAIC > 2), providing no evidence of sex-dependent IGE.

Given the evidence of sex-dependent IGE on FST phenotypes in CBA mice, we tested for IGE separately in males and in females. The multiple testing-corrected significance threshold we chose for the FST phenotypes accounted for our testing two FST phenotypes in two sexes, hence was p < 0.0125. The threshold for wound healing was p < 0.025, accounting for our testing in the two sexes independently. We found significant IGE for immobility in the first two minutes of the FST in females (two-sided t-test, t-stat = -3.34, p = 1.5e-3) but not in males (Fig. 3), consistent with the sex by genotype interaction predicted by the previous model comparison analysis. The other predictions of sex by genotype interactions from that analysis did not translate to significant genotype effects.

**Figure 3.**
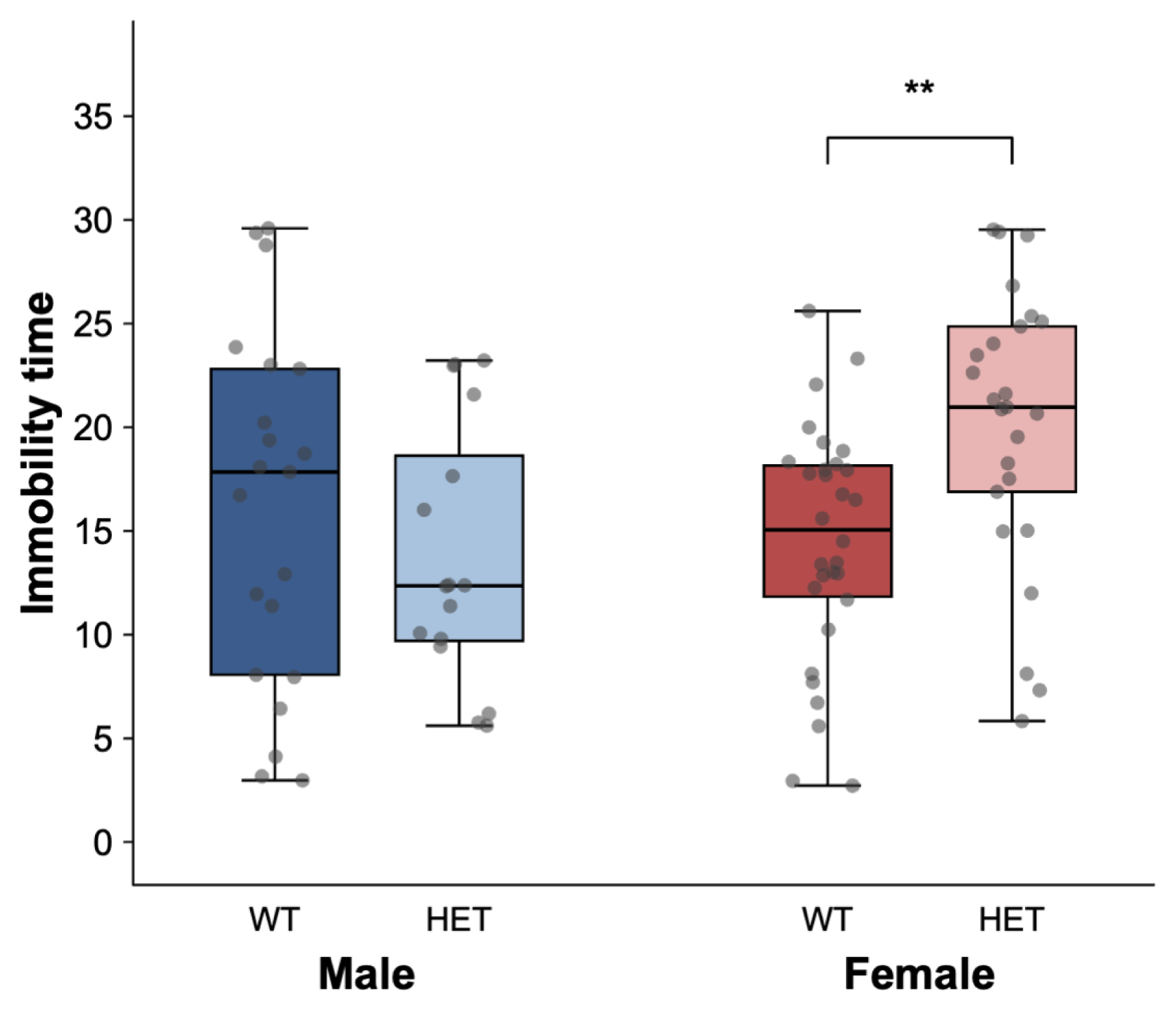
Female-specific IGE from *Epha4* in CBA/J focal mice. The phenotype, immobility during the first two minutes of the FST, was measured in CBA/J focal mice. Their cage mate was either *Epha4^+/+^* (WT) or *Epha4^+/-^*(HET). The multiple testing-corrected significance threshold we chose accounted for our testing two FST phenotypes in two sexes, hence was p < 0.0125.

### Global analysis of sex and strain as modulators of IGE from *Epha4*

The prior results (7) and the results presented above suggest that the strain of the focal mouse and the sex of the pair might modulate IGE from *Epha4*. To formally evaluate this possibility, we carried out a model selection analysis using the whole dataset (all four focal strains analysed jointly) and testing the explanatory variables focal strain, sex and cage mate *Epha4* genotype, as well as their interactions. We also included two additional models to test whether the similarity of the genetic backgrounds of the focal mouse and its cage mate was a better explanatory factor than the focal strain. It should be noted however, that the similarity model is the same as (fully confounded with) a model of B6-specific modulation of IGE (see caption of Fig. 2 and Methods). The models missing either sex or focal strain yielded very high AIC, indicating that both covariates are important factors influencing all three phenotypes considered (Fig. 4 and Supplementary Material, Table S6). For immobility during the first two minutes of the FST, there were five models that were competitive (ΔAIC < 2 (46)), including two models with interactions between sex and cage mate *Epha4* genotype and one with an interaction between focal strain and cage mate *Epha4* genotype. For immobility during the last four minutes of the FST, there were two competitive models, none of them with interactions with cage mate *Epha4* genotype. Finally, for wound healing, there were three competitive models, including one with an interaction between sex and cage mate genotype. Furthermore, the models testing for similarity-based epistasis were systematically better than the corresponding models of values-based epistasis. These results are consistent with, but do not demonstrate, modulation of IGE from *Epha4* by the similarity of the genetic backgrounds of the two mice in the cage and by sex, for stress-coping behaviour and wound healing. Notably, at least one model including cage mate *Epha4* genotype (i.e. IGE) was always among the competitive models, further supporting IGE from *Epha4* on these three phenotypes.

**Figure 4.**
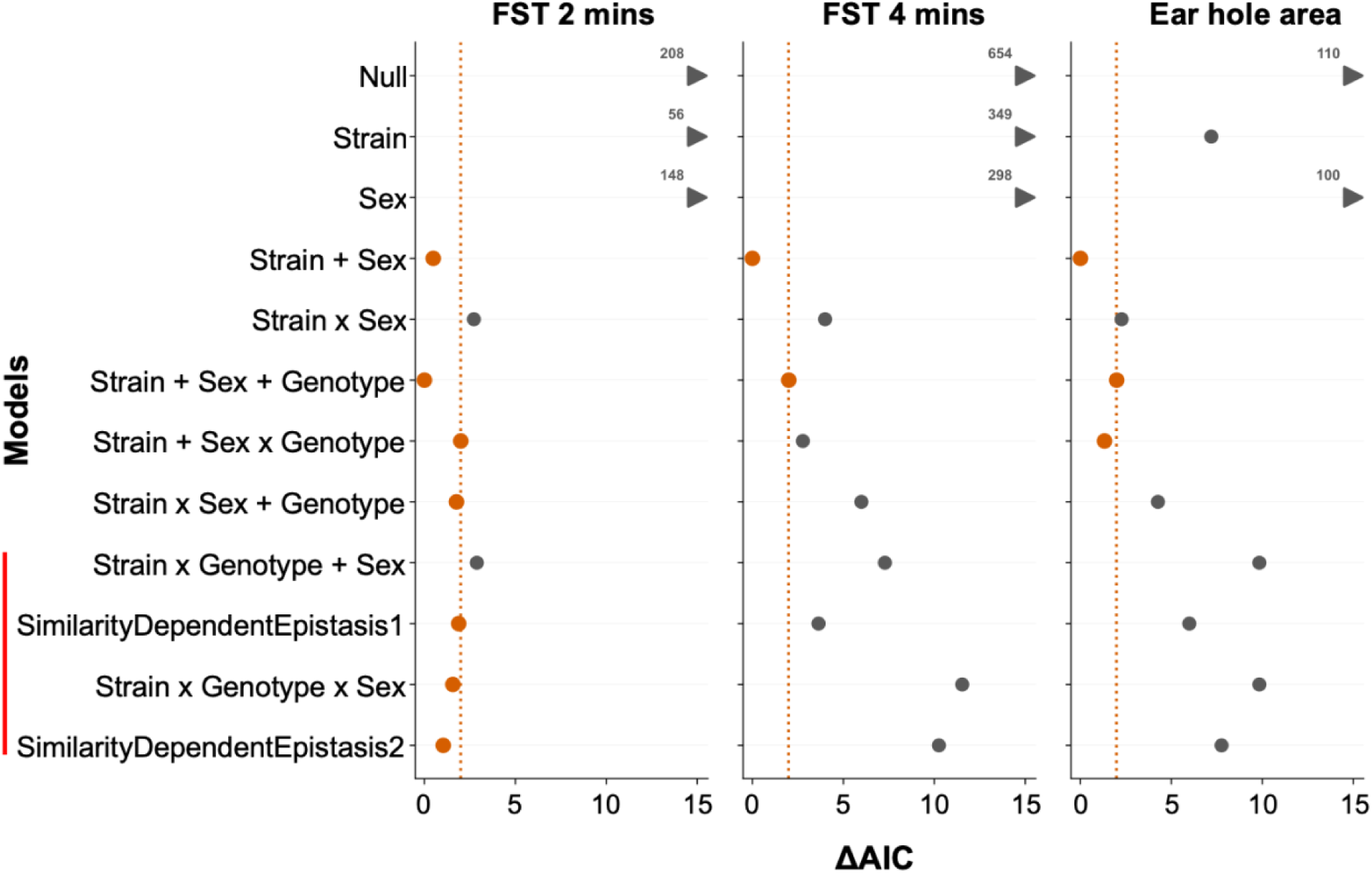
Evidence for social epistasis and modulation by sex. A model comparison analysis based on the AIC was carried out on all strains jointly. The best model is that with lowest AIC, and the ΔAIC in comparison with the best model is shown. Models within ΔAIC < 2 (to the left of the dotted line and coloured in orange) are competitive (46). The red bar highlights the four models including social epistasis. The models labelled “SimilarityDependentEpistasis1” and “SimilarityDependentEpistasis2” are shown underneath the corresponding models of “values-dependent social epistasis”.

### Deadly fighting is unlikely to have biased IGE analyses

Over the course of the experiment, one or both mice of 59 pairs got severe fight wounds and, as a result, died or had to be euthanised (“deadly fighting”). For those mice, no stress-coping behaviour or wound healing data were available, hence they were not included in the analysis of IGE from *Epha4* on these phenotypes. We investigated whether the presence/absence of severe fight wounds might have acted as a collider variable and induced spurious IGE from *Epha4* on stress-coping behaviour or wound healing. For collider bias to occur, the *Epha4* genotype would need to have affected the presence/absence of severe fight wounds and, independently, an unknown trait associated with immobility in the forced swim test or wound area would need to have also affected the presence/absence of severe fight wounds. We also investigated whether pleiotropic effects of *Epha4* on the presence/absence of severe fight wounds and on immobility in the forced swim test or wound area might have led to underestimated IGE (attenuation bias). To address both issues, we tested whether the Epha genotype of cage mates influenced the presence/absence of severe fight wounds.

We first observed that severe fight wounds (deadly fighting) occurred only in male cages, with 25% of male cages affected. In 92% of those cages, the focal mouse was aggressed. This could have suggested that B6 mice are very aggressive, but there was much less severe fighting in male cages with B6 focal mice (4% vs 74% respectively, two-sided Fisher’s exact test p = 5.8×10^-8^, Supplementary material, Table S7), suggesting that severe fighting occurred only when the genetic backgrounds of the two mice in the cage differed.

In male cages with mixed genetic backgrounds, we tested whether there was evidence of an effect of *Epha4* genotype on the presence/absence of severe fight wounds. There was no evidence of such an effect (two-sided Fisher’s exact test p > 0.05, Supplementary material, Table S8), making it unlikely that exclusion due to severe fight wounds created spurious IGE through collider bias or attenuated IGE through pleiotropic effects.

## Discussion

In this study, we characterised the context-dependency of IGE from *Epha4* on stress-coping strategy and wound healing. To be able to phenotype stress-coping behaviour in all 400 mice required for this study, we first developed and validated a new semi-automated pipeline to quantify immobility in the forced swim test from video recordings. Subsequently, we found several lines of evidence, including the results of a model selection analysis carried out on the joint dataset with all focal strains, that supported the modulation of IGE by the similarity of the genetic backgrounds and sex of the interacting mice. The most robust IGE were detected in CBA female mice for immobility during the first two minutes of the FST. We observed high levels of attrition due to severe fighting between male mice that had a different genetic background but found no evidence suggesting this had resulted in either spurious IGE (collider bias) or underestimated IGE (attenuation bias).

Social epistasis is an emergent property of focal and partner genotypes. In this study we investigated epistasis involving the *Epha4* genotype of cage mates and affecting stress-coping behaviour and ear wound healing. We investigated two possible architectures for social epistasis: “values-dependent” and “similarity-dependent”. The former was detected based on a statistical interaction with the strain of the focal mouse, whereas “similarity-dependent” epistasis was modelled based on the similarity of the genetic background of the focal mouse to the B6 genetic background of Epha4 cage mates. In our design, the variable “same genetic background as Epha4 cage mate” was fully confounded with “B6 is the focal strain”, therefore not allowing us to unequivocally implicate “similarity-dependent” epistasis. We nevertheless consider it likely, based on the “similarity-dependent” patterns of fighting we observed. Our reasoning is the following: first, our observations of severe fighting in many male cages, which were excluded from the analysis of IGE and epistasis, suggest that lower intensity aggression was common in those male cages that were included in the analysis. Secondly, aggression is unlikely to *mediate* IGE from *Epha4*, because severe fighting was not affected by the *Epha4* genotype, but that does not rule out a role as *modulator* of IGE arising from *Epha4*. Supporting this hypothesis, aggression has been shown to suppress allo-grooming (47), the trait of cage mates hypothesized by Baud et al. to mediate IGE from *Epha4* on both stress-coping strategy and ear wound healing (14). Aggression follows a similarity-dependent pattern, rather than a B6 background-dependent pattern, because the cage mates (B6 background) were almost always the aggressors yet there was very little severe fighting in the cages with a B6 focal mouse. The modulation of IGE from *Epha4* by aggression would therefore imply similarity-based social epistasis. To test this hypothesis and explore others, we have collected 24 hour-long videos for all the animals included in this study. However, the analysis of these videos requires the development, validation and application of advanced computer-vision models, hence is outside the scope of this report.

Complex scenarios underpinning IGE are increasingly being reported in the literature, including “developmental” IGE detectable after the source of the IGE is not present anymore (48), IGE “cascading” from one individual to another and then another one (49) and IGE being modulated by environmental factors (50,51). Social epistasis has also been considered before, but no general conceptual framework describing to the underlying processes has been proposed. We expand the prevailing view that social epistasis arises from differences in the responsiveness of the focal individuals associated with genotypes of the focal individuals (19,20) to potential modulation of that responsiveness by variants of social partners too and, in that case, either values-based or similarity-based epistasis, as well as potential modulation of the trait(s) of social partners mediating IGE (Fig. 5). Modulation at the level of the mediating trait(s) of social partners could involve increasing or decreasing the expression of one or more mediating traits, possibly in the extreme resulting in a change of mediating traits. Thus, all scenarios can change the magnitude of IGE, but only the latter will change the mechanisms of IGE. Importantly, similarity-based epistasis implies that two sets of variants in social partners act multiplicatively, and not additively as they would if two traits additively *mediated* IGE.

**Figure 5.**
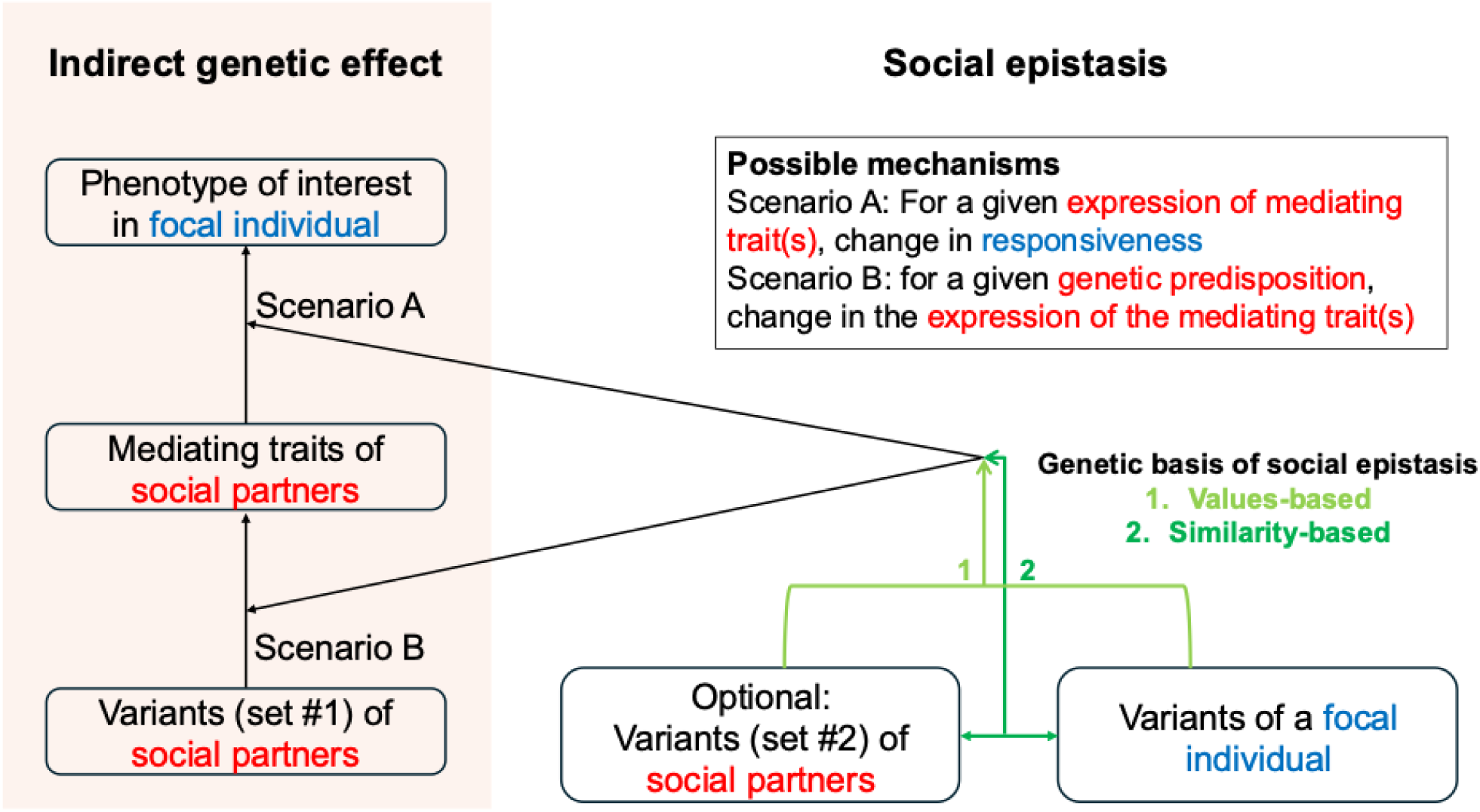
An expanded view of the processes underpinning social epistasis.

In addition to social epistasis, our results indicated modulation by sex. Because all groups were same-sex, as is usual with laboratory rodents to avoid mice uncontrollable breeding, the variable was sex of the dyad. Our results shed no light on what might occur in mixed groups of males and females. If aggression also underpins the modulation by sex, however, we expect mixed-sex dyads to show little to no aggression, hence behave like cages of females, rather than like cages of males. As for social epistasis, the modulation of IGE by sex could occur at the level of the responsiveness of the focal individuals or at the level of the expression of the traits of social partners mediating IGE.

Floor and ceiling effects might explain some of the apparent interactive effects that we observed, when the phenotype in one category (sex or strain) is so high or so low that differences associated with the *Epha4* genotype of cage mates are difficult or impossible to observe. Baud et al. suspected this was an issue for FST immobility in FVB females (14) (floor effect), and in the present study was likely an issue for FST immobility in MRL mice, especially females (ceiling effect).

The context-dependency of IGE has strong implications for the design of experiments aimed at dissecting the mechanisms of IGE: effect sizes estimated in one context have limited portability across contexts (populations, strains, sexes, etc.), just as direct genotype–phenotype associations do (16). In the case of IGE from *Epha4*, high prospective power to detect an effect previously observed in FVB did not translate to detection in all contexts studied here. CBA females were the only context in which IGE from *Epha4* were significant and are the evidence-based context of choice for future studies aimed at understanding the neurobehavioural and physiological mechanisms underlying IGE from *Epha4*.

An important contribution of our work is the development of a semi-automated pipeline for converting EthoVision pixel intensity-based activity measurements into validated mobile/immobile calls for each frame of an FST video recorded from the side. Only 6–7 manually annotated videos per strain were required to calibrate the activity threshold, substantially reducing the manual-scoring burden relative to annotating the entire dataset. Our pipeline starts off from an output from EthoVision, a proprietary (but widely used) software, but it could easily be adapted to take as input the same output from any other software.

Our experimental design implied various sources of visual heterogeneity potentially affecting such a pipeline, some of which had a genetic basis (i.e. strain differences): four different coat colours (ranging from white MRL mice to black B6 mice), two different background colours (black and white) used to maximise contrast with the mouse’s coat colour, different body sizes (ranging from large MRL mice to small B6 or DBA mice). In addition, technical variation arose from an unplanned change in lighting, creating a glare in some of the videos, and slightly different zoom values incurred by switching on and off the camera between test days. We demonstrated that our pipeline effectively accounts for this heterogeneity, ensuring high-performance levels across strains in particular (Figure 2D).

## Methods

### Power analysis carried out prior to the experiment

A G*Power analysis based on the effect size (mean and standard deviation) of IGE from *Epha4* on stress-coping behaviour in male FVB mice of the experiment reported by Baud et al. determined that 98 mice per focal strain were required to achieve 95% power to detect IGE from *Epha4* on that phenotype (comparison of 49 focal mice paired with a wild-type *Epha4^+/+^* cage mate and 49 focal mice paired with a heterozygous *Epha4^+/-^*cage mate). That sample size was expected to give 97% power to detect IGE on the wound healing phenotype.

### Animals

All experiments followed the principle of the “3Rs” (replacement, reduction and refinement) according to European Union Directive 63/2010 and its implementation in the Member States. The study was conducted according to the guidelines of the local law (32/2007) and European regulations (2010/63/EU) and approved by the Ethics Committee of Parc de Recerca Biomèdica (Comité Ético de Experimentación Animal del PRBB (CEEA-PRBB); AB 11687). The CRG is authorized to work with genetically modified organisms (A/ES/05/I-13 and A/ES/05/14).

The focal strains C57BL/6J (“B6”), CBA/J (“CBA”), DBA/2J (“DBA”) and MRL/MpJ (“MRL) were purchased from the Jackson Laboratory (JAX) then bred at the PRBB Parc de Recerca Biomèdica de Barcelona Animal Facility in Barcelona, once at the beginning of the experiment and once halfway through the experiment, to avoid genetic drift. The Epha4<tm1Byd> (Epha4) cage mates were imported from the laboratory of Dr. Abraham A. Palmer (University of California San Diego, CA, USA), from the same colony that was used for Baud et al. (14). This knockout model was originally created by (52), then donated to Dr. Elena Pasquale (Sanford Burnham Prebys, San Diego, CA, USA), who then donated it to Dr Baud. A full characterization of the genetic background of the Epha4 line by Transnetyx (Transnetyx Genotyping Services, Cordova, TN) using the MiniMUGA genotyping array showed that it had a mixed background, primarily B6 but also, as expected from how the model was created, C57BL/10 and 129S2. To reduce background heterogeneity and refresh the Y, X and mitochondrial chromosomes, upon arrival in Barcelona the Epha4 line was backcrossed to B6 male and female mice for three generations. Throughout the experiment, to generate *Epha4^+/-^*cage mates *Epha4^+/-^*mice were bred with B6 mice, further homogenising the genetic background of the Epha4 mice used in this experiment. Sperm from *Epha4^+/-^*mice was cryopreserved at the end of the experiment and is available from Charles River France upon request to Amelie Baud.

Mice were housed in a specific pathogen free (SPF) area of the Parc de Recerca Biomèdica de Barcelona (PRBB) animal facility under controlled conditions: 22°C ± 2°C, 40%–70% humidity, a 12-hour light cycle, and *ad libitum* access to food and water.

### Genotyping

Tails tips were collected at the end of the experiment from Epha4 cage mates and sent to Transnetyx Inc. (Transnetyx Genotyping Services, Cordova, TN) for genotyping. Primers used were: Fw-WT: ACCGTTCGAAATCTAGCCCAG and RV-WT ACCGTTCGAAATCTAGCCCAG with a product size of 365 bp indicating the presence of an *Epha4* WT allele; Fw-KO GCTTCCTCGTGCTTTACGGTATC and RV-KO AGCCTTGCCACCTGGAGC, with a product size of 550 bp indicating an excised (i.e. KO) *Epha4* allele.

### Experimental design

Focal mice from the four inbred strains (B6, DBA, MRL or CBA) were paired with *Epha4*^+/+^ (WT) or *Epha4*^+/-^(Het) cage mates at weaning, prior to the establishment of dominance hierarchies, in same-sex pairs. About half the cages were males, half females. The pairing was done on a rolling basis rather than as a few large batches. The experimenter was blind to the genotype of the Epha4 cage mate throughout the experiment (no difference in coat colour and genotyping done at the end of the experiment). Immediately prior to pairing, the focal mice were punched in both ears (wound healing test, see below).

After pairing, the mice were left undisturbed for six weeks, during which they interacted freely in the cage. A special monitoring protocol was implemented to ensure animal welfare and minimize experimenter-induced perturbations and stress. Health checks were done without opening the cage unless pre-defined welfare concerns were detected, in which case the cage was opened, and additional observations were made.

At about nine weeks of age, the pairs spent 24 hours in an enclosure with bedding, food and water to collect videos for scoring additional behaviours (results not reported in this study). About three days later, the focal mice were tested in the forced swim test assay, to characterize their stress-coping behaviour (see below). Immediately after, both the focal mouse and its Epha4 cage mate were euthanized, and the ears of the focal mouse collected for wound healing characterisation (see below)

The final sample sizes were 107 B6, 97 CBA, 101 DBA, and 95 MRL mice paired in roughly equal proportions with *Epha4+/-* and *Epha4+/+* cage mates. An additional 59 pairs were euthanised due to severe fighting wounds, following the strict supervision and decision protocol that had been approved.

### Estimating the rate of ear wound healing

Both ear pinnae of the focal mice were punched at weaning using 2 mm (diameter) ear punch scissors (Scientific Hub Services, #KN-292-2-2). Both ears were collected after euthanasia and stored in 10% buffered formalin phosphate (Electron Microscopy Sciences, #15741-01) at room temperature until analysis. To measure the area of the ear hole after weeks of healing from the ear punch, each ear was mounted on a histology slide, and photos were taken from a fixed distance. Images were analyzed with the ImageJ software (53) to delimitate and measure the hole area. Ear holes that were open from the time of punching were excluded from the analysis, but ear holes that opened due to the healing process were included.

### Forced Swim Test (FST)

At about 9.5 weeks of age the focal mice were tested in the FST. In the morning (light phase of the light-dark cycle), mice were brought to the testing room for at least 15 minutes for habituation and then the focal mice were placed in 30 cm height x 20 cm diameter glass buckets filled with water at 23-26°C. The water was changed between each mouse. Mice were video recorded from the side against either a white (for B6, CBA and DBA mice) or a black (for MRL mice) background, to ensure maximum contrast. The zoom of the camera was adjusted once per video recording session (day), which was accounted for in the analysis of the videos (see below).

### Immobility time analysis (Fig. 2)

#### 1. Processing with EthoVision

The videos were analyzed using the animal tracking system EthoVision XT 16 (Noldus), ‘Activity analysis’ mode. The software requires the definition of an “arena” that will be the focus of the analysis: for each new video, the scale (pixel to cm) was set using the width of the water bucket, which was 15cm, then the arena was defined with a fixed width and height (in cm) that encompassed the full width of the water bucket and all the vertical space where mouse or water movement could occur. We then took note of the width and height of the arena in pixels for future normalization of zoom. The trial control settings used a logic of starting the trial as soon as the center point of the animal was detected in the arena for more than 1s. Next, the “activity threshold” and “background noise threshold” were both set to 5 for all videos. Along with this the ‘Compression artifacts filter’ was switched on. These settings were chosen by comparing, for a few videos, the rendering of movement given by EthoVision (displayed in purple on the video) to mouse movements, avoiding the capture of water movements. With these thresholds set, for every frame in a video the software returns the proportion of pixels affected by mouse movement (“Activity%”).

All Activity% values in a video were corrected with (divided by) a “Zoom Value” (see below) to account for the zoom of the camera used for videorecording. Specifically:

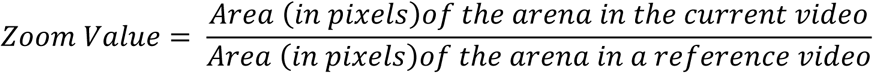

After doing so, we also smoothed the activity values by replacing the value of activity for each frame by the average activity value for the 100 frames around it.

#### 2. Learning Activity% thresholds

Then, strain-specific thresholds were determined for converting the smoothed activity values to a binary call of mobile or immobile for each frame. For MRL focal mice, lighting differences were observed for videos taken after a certain date. Hence two batches of MRL videos were identified (before and after “experiment 199”) and individual thresholds estimated, tested and applied separately for the two batches. The testing and evaluation results are, however, shown as one in Fig. 2D and Supplementary Material, Fig. S1. The thresholds were determined using a training set consisting of all 6 minutes of 7-10 videos from each of the four focal strains. These videos were annotated by an experimenter making a call of mobile or immobile every 5s (54–57). To compare machine-made and experimenter-made calls, each experimenter mobile/immobile call was extended to the 50 frames corresponding to 1s on either side of the frame annotated by the experimenter. The thresholds were then determined by maximizing the average F-1 score (the experimenter’s calls are treated as the ground-truth here) defined as:

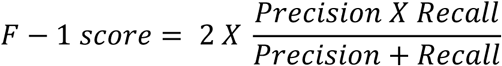

with

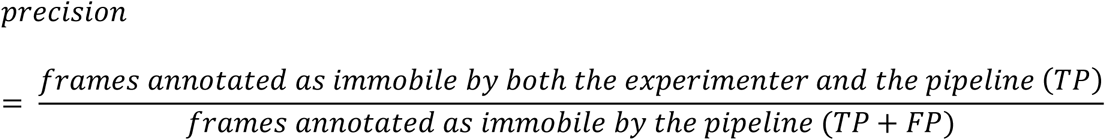

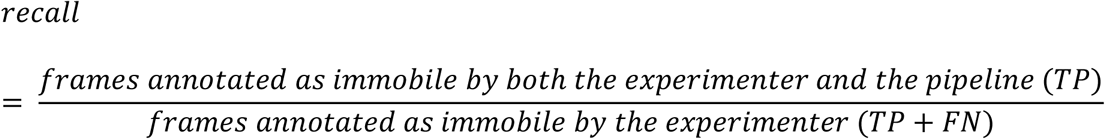

#### 3. Evaluating the automated pipeline against manual annotations

To evaluate the whole pipeline, all 6 minutes of a subset of 3-4 videos per focal strain was used (test set, different from the training set). As with the videos of the training set, these videos of the test set were annotated by the experimenter every 5s; mobile/immobile calls extended to all frames within 1s on either side and an F-1 score was calculated per video and averaged across all videos in the test set to produce Fig. 2D.

#### 4. Applying the pipeline

After finalising the pipeline, all data (including those used in the training and test sets) were used for statistical analysis of genetic effects.

### Statistical analysis

Three phenotypes were investigated in this study:

a. Immobility during the first 2 minutes of the FST (a measure of stress-coping behaviour)
b. Immobility during the last 4 minutes of the FST (a measure of stress-coping behaviour)
c. Ear hole area (a measure of wound healing)

All analyses were carried out in Python. The software versions used are as follows:

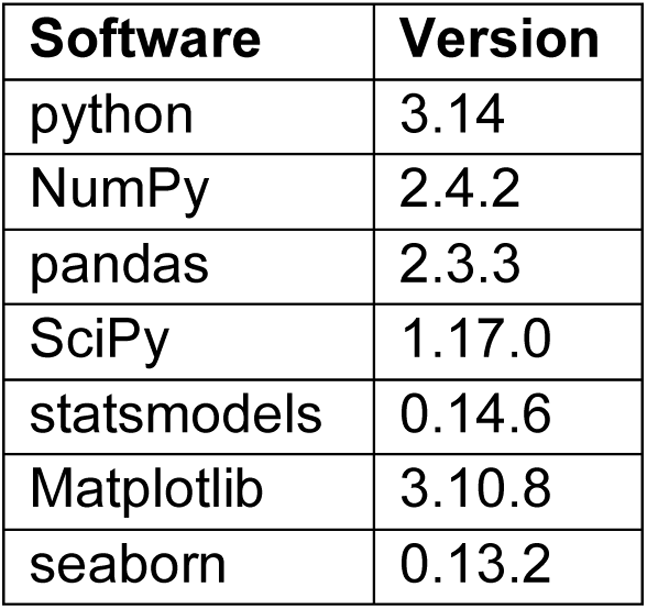

| Software | Version |
| --- | --- |
| python | 3.14 |
| NumPy | 2.4.2 |
| pandas | 2.3.3 |
| SciPy | 1.17.0 |
| statsmodels | 0.14.6 |
| Matplotlib | 3.10.8 |
| seaborn | 0.13.2 |

#### Normalization

Prior to fitting linear models, the phenotypes were normalized using the Box-Cox power transformation (58)(scipy.stats.boxcox function). Each phenotype was normalized using all samples regardless of strain or genotype. For the transformation to work, all 0 values were increased to 0.01 regardless of the phenotype. λ values that were picked by the function for the transformation were as follows:

| Phenotype | $\lambda$ values |
| --- | --- |
| FST immobility – first two minutes | 0.7737976617218821 |
| FST immobility – last four minutes | 4.262909867373311 |
| Ear hole area | 0.53207656449072 |

#### Linear models

The contribution of focal strain (genetic background), sex, *Epha4* cage mate genotype, and all possible interactions of these variables to phenotypic variation was evaluated using linear models for the two FST phenotypes (statsmodels.regression.linear_model.OLS function). For ear hole area, a mixed model was used (statsmodels.regression.mixed_linear_model.MixedLM function) to account for the inclusion of two data points (ears) per mouse.

Nested linear models were compared using analysis of variance (statsmodels.stats.anova.anova_lm). For mixed-effect models, a likelihood ratio test was used (using statsmodels.regression.linear_model.RegressionResults.llf and scipy.stats.chi2).

To compare multiple models, including non-nested models, the Akaike Information Criterion (AIC) was used (statsmodels.tools.eval_measures.aic function).

## Data availability statement

All the raw data files for the phenotypes (FST videos and ear pictures), along with phenotype data, the necessary metadata of the animal colony (ID, sex, strain, genotypes) and the metadata for running the automated FST method are available at https://doi.org/10.5281/zenodo.23017774. Sample data, the automated FST method and the code used for IGE analysis are available at https://github.com/Baud-lab/Epha4_IGE_CRG/tree/main

## Acknowledgements

This work was supported by PID2021-122651NA-I00 and PRE2022-101263 contracts funded by MCIN/AEI/10.13039/501100011033 and FSE+ to AB. We acknowledge support of the Spanish Ministry of Science and Innovation through the Centro de Excelencia Severo Ochoa (CEX2020-001049-S, MCIN/AEI /10.13039/501100011033), the Generalitat de Catalunya through the CERCA programme and to the EMBL partnership. We are grateful to the staff of the PRBB animal facility for their support. We acknowledge client support from Noldus. During the preparation of this manuscript, the authors used the family of ChatGPT-5.x models for exploring the scientific literature relevant to the topic of this manuscript and assistance in writing and debugging Python scripts. In all cases, the outputs were critically reviewed and manually edited by the authors, who take full responsibility for the scientific content of the final manuscript.

## Authors’ contributions

A.B. designed the study and acquired funding. P.M, M.S. and S.N. carried out the mouse experiment. P.M. analysed the data. A.B. and P.M. wrote the manuscript. All authors reviewed and approved the final draft.

